# Efficient target cleavage by Type V Cas12a effector programmed with split CRISPR RNA

**DOI:** 10.1101/2020.12.21.423781

**Authors:** Regina Tkach, Natalia Nikitchina, Nikita Shebanov, Vladimir Mekler, Egor Ulashchik, Olga Sharko, Vadim Shmanai, Ivan Tarassov, Konstantin Severinov, Nina Entelis, Ilya Mazunin

## Abstract

CRISPR RNAs (crRNAs) directing target DNA cleavage by type V-A Cas12a nucleases consist of repeat-derived 5’-scaffold moiety and 3’-spacer moiety. We demonstrate that removal of most of the 20-nucleotide scaffold has only a slight effect on *in vitro* target DNA cleavage by Cas12a ortholog from Acidaminococcus sp (AsCas12a). In fact, residual cleavage was observed even in the presence of a 20-nucleotide crRNA spacer part only, while crRNAs split into two individual moieties (scaffold and spacer RNAs) catalyzed highly specific and efficient cleavage of target DNA. Our data also indicate that AsCas12a combined with split crRNA forms a stable complex with the target. These observations were also confirmed in lysates of human cells expressing AsCas12a. The ability of the AsCas12a nuclease to be programmed with split crRNAs opens new lines of inquiry into the mechanisms of target recognition and cleavage and will further facilitate genome editing techniques based on Cas12a nucleases.

## INTRODUCTION

CRISPR (clustered regularly interspaced short palindromic repeats) and CRISPR-associated proteins (Cas) constitute adaptive immune systems of bacteria and archaea (1). These systems act through CRISPR RNA (crRNA)-dependent targeting of foreign nucleic acids by Cas nucleases. At present, two classes, six types and numerous subtypes of CRISPR-Cas systems have been described and some have found wide use in genomic editing. AsCas12a, a class 2 Type V-A Cas12a effector nuclease from Acidaminococcus sp., has been applied for genome editing in a number of organisms, including human cells (2, 3, 4). The crRNA of Cas12a effector consists of two nearly equally sized moieties, a repeat-derived scaffold (also referred to as the 5’-handle) and a spacer or guide segment complementary to target DNA sequences. To date, structures of three type V-A Cas effector proteins, AsCas12a (5), LbCas12a (6), and FnCas12a (7) charged with crRNA and bound to target DNA have been determined. These structures revealed that crRNA scaffold adopts a pseudoknot structure rather than a simple stem-loop fold predicted from its nucleotide sequence (5). Cas12a proteins interact with this pseudoknot (6) and recognize T-rich protospacer-adjacent motifs in target DNA. The Cas12a/ crRNA complex induces a staggered double-stranded break in the target region complementary to crRNA spacer part (5, 8).

Previous studies analysed crRNA structure requirements for DNA cleavage by Cas12a nucleases (6, 7, 9). For example, it was demonstrated that extending the length of crRNA at the 5′ end enhanced mammalian genes editing efficiency by AsCas12a (10). This approach was used to extend AsCas12a crRNAs with switchable domains activated by trigger RNAs via strand displacement, which can be adapted for the transcriptional control of gene expression in *<u>E</u>*. *<u>coli</u>* (11). On the other hand, shortening of the scaffold and/or spacer crRNA domains decreased FnCas12a and LbCas12a cleavage activities (8, 9). It was also shown that the FnCas12a nuclease activity depends on the secondary structure of crRNA scaffold and is sensitive to nucleotide changes in this region (8).

Here, we report the data on the effects of crRNA truncation on the *in vitro* DNA cleavage by the AsCas12a nuclease (2, 3, 4). We demonstrate that the cleavage activity persists after substantial reduction of the crRNA scaffold length and can even be detected upon complete removal of the scaffold, i.e., in the presence of a 20-nucleotide RNA matching the spacer part only. Further, we show that the residual nuclease activity of AsCas12a charged with spacer RNA can be restored by the *in trans* addition of a 20-base scaffold RNA. In a pure *in vitro* system and in human cell lysates the efficiency of cleavage by Cas12a effector charged with split crRNA is comparable to that observed with intact crRNA. The use of split crRNAs composed of the constant scaffold part and a variable spacer parts should further simplify genome editing by Cas12a effectors and may allow new ways to fine-tune the editing process.

## MATERIAL AND METHODS

### Expression and purification of AsCas12a and SpCas9 nucleases

For the subsequent testing of various crRNA/sgRNA variants in an *in vitro* cleavage assay, AsCas12a and SpCas9 nucleases were expressed from the pET-based T7 promoter-containing plasmids (Addgene, Plasmids #90095, #62374) in Escherichia coli strain BL21-DE3 as described (3) with modifications. Briefly, the lysate of cells was loaded onto a HiTrap Chelating HP column (GE Healthcare) equilibrated with a buffer containing 20 mM imidazole. The recombinant protein was eluted with a buffer containing 500 mM imidazole, mixed with TEV protease, and dialyzed at 4°C for 12 hours. For SpCas9 nuclease, the TEV protease cleavage step was omitted. Then the proteins were concentrated with Amicon Ultra 10K filter (viMillipore) and applied to HiLoad Superdex 200 16/60 column (GE Healthcare). Fractions containing either AsCas12a or SpCas9 were collected, concentrated and then snap-frozen in liquid nitrogen in small aliquots that were stored at −80°C until use. The protein concentration was measured with Qubit Protein Assay Kit (Thermo Fisher) according to the manufacturer’s instructions. The protein quality was tested by SDS-PAGE gel electrophoresis and visualized after Coomassie blue staining by Gel Doc EZ Imager (Bio-Rad).

### *In vitro* cleavage assay with purified proteins

*In vitro* cleavage assay with the purified AsCas12a and SpCas9 nucleases was carried out in a volume of 30 µL in 1× NEB2.1 buffer (New England Biolabs Inc.). The standard reaction containing 10 nM of PCR-obtained DNA template, 500 nM of the nuclease, and 5000 nM of crRNA were incubated 30 min at 37°C; the reaction was stopped by the addition of 40 µg/µL of Proteinase K and incubated at 37°C for 15 min. Analysis of the cleavage was performed by electrophoresis on a 1% agarose gel in 1× TBE buffer. EtBr staining gels were visualized by Gel Doc EZ Imager (Bio-Rad) and quantified with Image Lab™ Software 6.0.1 (Bio-Rad).

The oligoribonucleotides synthesis and characterization are described in Supplementary Data. DNA substrates for *in vitro* cleavage represent fragments of human mitochondrial DNA amplified by PCR in Q5® High-Fidelity 2X Master Mix (M0492L, NEB). As a template, genomic DNA isolated from T-REx cell line (T-REx™-293 Cell Line, Invitrogen, kat. # R71007) by GeneJET Genomic DNA Purification Kit (K0722, Thermo Scientific™) was used. DNA concentration was measured by Qubit 3.0 fluorometer using Qubit DNA BR Assay Kit (Q32853, Thermo Scientific™). Oligonucleotide primers provided by Lumiprobe RUS Ltd are listed in Table S1 SD.

### Fluorometric measurements

Fluorescence measurements were performed using a QuantaMaster QM4 spectrofluorometer (PTI) in binding buffer (20 mM Tris HCl (pH 7.5), 50 mM NaCl, 0.1 mM DTT and 5 mM MgCl2) at 25°C. Unmodified and chromophore-labeled DNA oligonucleotides, AsCas12a protein, crRNA and its scaffold and spacer fragments used in fluorometric experiments were purchased from Integrated DNA Technologies. Double-stranded DNAs and beacon DNA were formed by mixing equimolar amounts of synthetic complementary strands (final concentrations were within low µM range) in a buffer containing 20 mM Tris, pH 7.5, 100 mM NaCl; heating for 2 min at 90°C and slowly cooling the reactions to 20°C. The beacon DNA construct shown in Figure 7B was previously described in (12). The protospacer adjacent motif (PAM)-distal ends of the beacon target and non-target strands are labeled with fluorescein and Iowa BlackFQ, respectively. If not otherwise stated, final assay mixtures (800 µl) contained 40 nM of AsCas12a protein, 1 nM Cas beacon and crRNA or its fragments (scaffold and spacer) at various concentrations. The fluorescein fluorescence intensities were recorded with an excitation wavelength of 498 nm and an emission wavelength of 520 nm. Time-dependent fluorescence changes were monitored after addition of negligible volume of Cas beacon to a cuvette followed by manual mixing; the mixing dead-time was 15 s.

### *In vitro* cleavage assay in hAsCas12a cell lysate

Using the Flp-In™ System, we introduced the hAsCas12a gene into the human HEK 293 T-REx cell line nuclear genome (Flp-In ™ T-REx ™ −293 Cell Line, Invitrogen) allowing controlled expression by the tet-on system. Successful integration was monitored by antibiotic selection with hygromycin B (100 µg ml−1, Invivogen) and blasticidin (5 µg ml−1, Invivogen). Expression of AsCas12a nuclease in the stable HEK 293 T-REx cell line was induced by 100 ng/mL tetracycline for 24 h and assayed by Western immunoblotting using FLAG-tag specific antibodies (Sigma-Aldrich, F1804).

Cells with inducible expression of the AsCas12a nuclease were cultured at 37°C, 5% CO2 in standard essential modified Eagle’s medium (EMEM) containing 4.5 g/L glucose, 1 mM pyruvate, and 5 mg/mL uridine supplemented with 10% fetal bovine serum. The expression of the nuclease was activated at 60-80% confluency by the addition of tetracycline to a final concentration of 100 ng/mL. 24 hours post induction, cell lysate was prepared and *in vitro* cleavage reactions were carried out as described (13) at DNA:RNA ratio of 1:30. Analysis of the cleavage was performed by electrophoresis on a 1% agarose gel in 1×TAE buffer. EtBr staining gels were visualized by Hero Lab Transilluminator UVT-28M (Herolab GmbH) and quantified with Image Lab™ Software 6.0.1 (Bio-Rad).

## RESULTS

### The AsCas12a nuclease can cleave DNA targets when programmed with 5’ truncated crRNAs

We performed an *in vitro* DNA cleavage assay using purified AsCas12a nuclease and a set of chemically synthesized crRNAs molecules. We tested crRNAs truncated from the 5’-end, i.e., harbouring scaffold parts of different length (versions −17, −15, −7 and −2, Figure 1A) and a constant 20-nt spacer part. An RNA that consisted of a scaffold extended by just two extra bases of the spacer (version +2) was also tested. As a DNA cleavage substrate, a PCR fragment containing a protospacer fully matching crRNA spacer with a functional PAM was used. The AsCas12a nuclease was surprisingly tolerant to 5’-truncations of the scaffold. As can be seen from Figures 1A and 1B, at the conditions of the experiment more than 80% of the DNA substrate was cleaved when the crRNA scaffold part was reduced to just seven nucleotides (version −7). The −2 version, as well as the +2 crRNA (a full scaffold with 2 spacer nucleotides at the 3’-terminus) were inactive.

**Figure 1.**
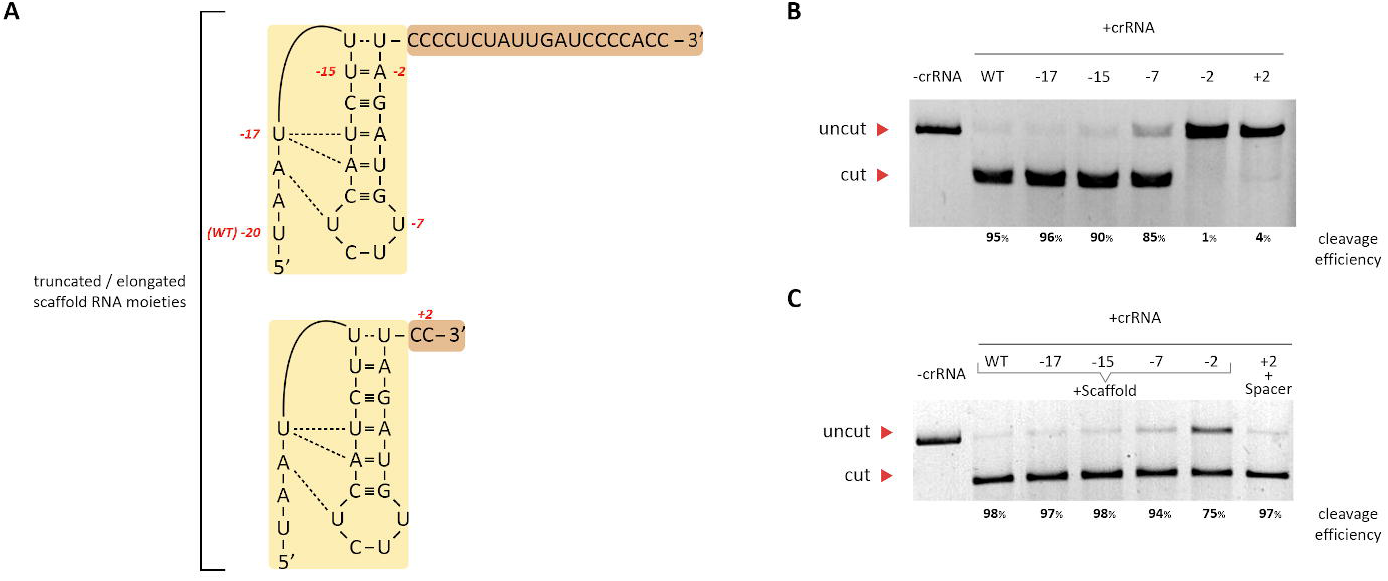
Effect of crRNA scaffold length on AsCas12a cleavage activity. **(A)** The sequence of the crRNA used in the study; the structure of the scaffold region is shown as in (5); the numbers in red indicate the 5’-terminal nucleotides of truncated forms. In the +2 version shown below, 18 nucleotides from the 3’-part of the spacer were eliminated. The scaffold part is coloured yellow, the spacer is brown. **(B)** The products of *in vitro* cleavage of DNA template (“-crRNA”) by AsCas12a in the presence of indicated crRNAs. The reactions contained 10 nM of PCR-amplified target DNA fragments, 500 nM of recombinant AsCas12a, and 5000 nM of crRNAs (a 1:50:500 ratio), and was incubated 30 min at 37°C. **(C)** Truncated crRNAs forms were supplemented with equimolar amounts of full-sized scaffold RNA, combined with AsCas12a and the DNA target and cleavage reaction was performed. A reaction containing +2 scaffold was supplemented with 20-nt RNA matching the spacer. Reaction products were separated in 1% agarose gels and stained with ethidium bromide. Here and in the following figures, a representative image (from three independent repeats) is shown. The cleavage efficiencies (calculated as a ratio of intensities in cut and uncut DNA bands) are shown below each gel.

We next repeated the experiment by combining AsCas12a, various crRNAs and a 20-nt RNA corresponding to the complete scaffold part of intact crRNA. As can be seen from Figure 1C, scaffold RNA added *in trans* stimulated target cleavage in the presence of the −2 crRNA version (75% of target cleaved compared to ∼2% cleavage obtained with the −2 crRNA alone). The extended, +2 version of the scaffold had an even stronger stimulatory effect on cleavage programmed by the −2 crRNA (97% of target cleaved). At the conditions of the experiment, neither spacer nor scaffold RNA alone induced target cleavage.

We next investigated the kinetics and concentration dependence of target DNA cleavage by AsCas12a charged by either full-sized or truncated (version −7) crRNAs. As can be seen from Figure 2A, in reactions containing full-sized crRNA ∼70% of target DNA was cleaved just after 1-minute incubation at 1:5:50 and 1:50:500 target DNA:AsCas12a:crRNA ratios. In reactions containing truncated crRNA, comparable efficiency was reached only after 5 min of the reaction at 1:50:500 DNA-AsCas12a-crRNA ratio (Figure 2B). At the 1:5:50 DNA:AsCas12a:crRNA ratio cleavage was low, suggesting that the −7 crRNA may be defective in binding to the effector.

**Figure 2.**
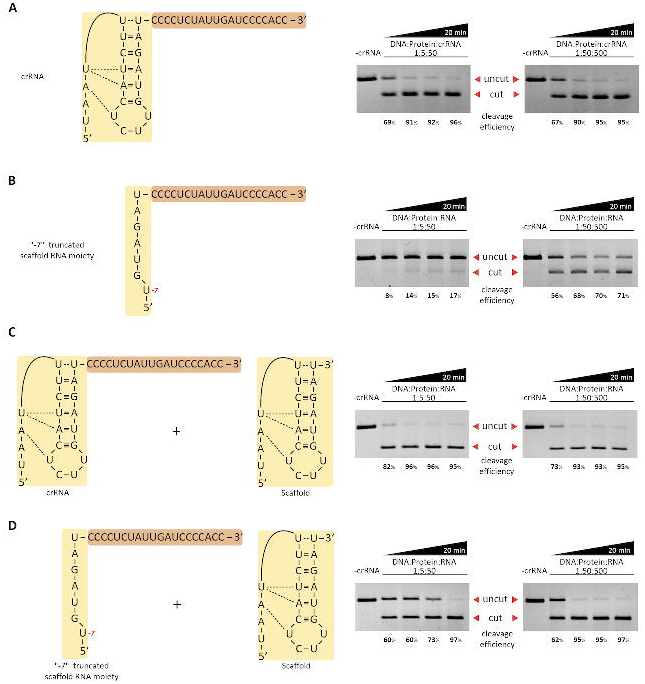
Effects of incubation time and concentration of reaction components on AsCas12a cleavage activity. **(A)** The products of *in vitro* cleavage of target DNA (“-crRNA”) by AsCas12a in the presence of full-sized crRNA at four times points (1, 5, 10, and 20 minutes of reaction) at two different DNA:AsCas12a:crRNA ratios, 1:5:50 (left) and 1:50:500 (right). **(B)** As in (A), but using −7 crRNA. **(C)** As in (A), but using full-sized crRNA and equimolar amounts of scaffold RNA. **(D)** as in (A), but using - 7 crRNA and equimolar amounts of scaffold RNA.

### The *in trans* addition of scaffold RNA stimulates target DNA cleavage by AsCas12a programmed with 5’ truncated crRNA

When cleavage reactions were performed using intact crRNA and complete scaffold part added *in trans*, little effect was observed compared to reactions containing just intact crRNA (Figure 2C). How-ever, when the truncated −7 version of crRNA was combined with the scaffold part added *in trans*, efficient cleavage was observed even at low DNA:AsCas12a:crRNA ratio, when the −7 version alone was poorly active (compare Figure 2B and Figure 2D).

We next asked whether AsCas12a charged with two 20-nt RNAs corresponding to complete spacer and scaffold domains of intact crRNA will be able to induce specific cleavage of the DNA substrate. As can be seen from Figure 3, the simultaneous addition of two crRNA domains (a condition further referred to as “split crRNA”) induced specific cleavage of the target, which was only slightly less efficient than that induced by full-sized crRNA (90% versus 96%) at the conditions of the experiment. Neither the spacer nor the scaffold RNAs taken separately induced appreciable substrate cleavage at these conditions.

**Figure 3.**
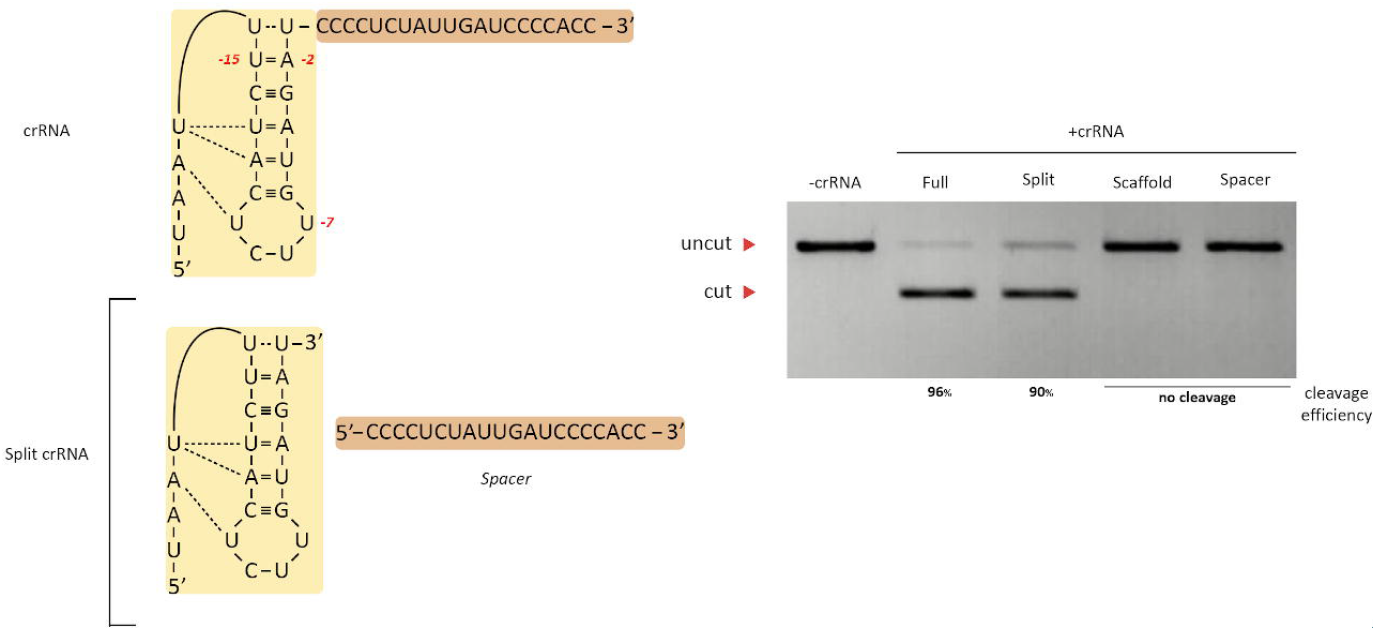
Split crRNA promotes efficient target cleavage by AsCas12a *in vitro*. A representative image (from three independent repeats) of an agarose gel with *in vitro* DNA cleavage reaction products formed in the presence of 3 µM RNAs (at a 1:30:300 DNA:AsCas12a:RNA ratio) is shown on the left. “Split” refers to reactions containing both scaffold and spacer RNAs.

To compare the binding of intact and split crRNA to AsCas12a, we performed *in vitro* DNA cleavage assay in the presence of equimolar amounts of another crRNA (hereafter referred to as “ crRNA competitor”) targeting an unrelated sequence not present in the DNA template. As can be seen from Figure 4A, the competitor had no effect on target DNA cleavage by effector charged with intact cognate crRNA but abolished cleavage by effector charged with split crRNA. The inhibition of target cleavage by effector charged with split crRNA by competitor crRNA was dose-dependent (Figure 4B) and became complete when the concentration of competitor crRNA reached 20% of split crRNA components concentration. We thus conclude that split crRNA components have reduced affinity for the AsCas12a nuclease compared to full-sized intact crRNA.

**Figure 4.**
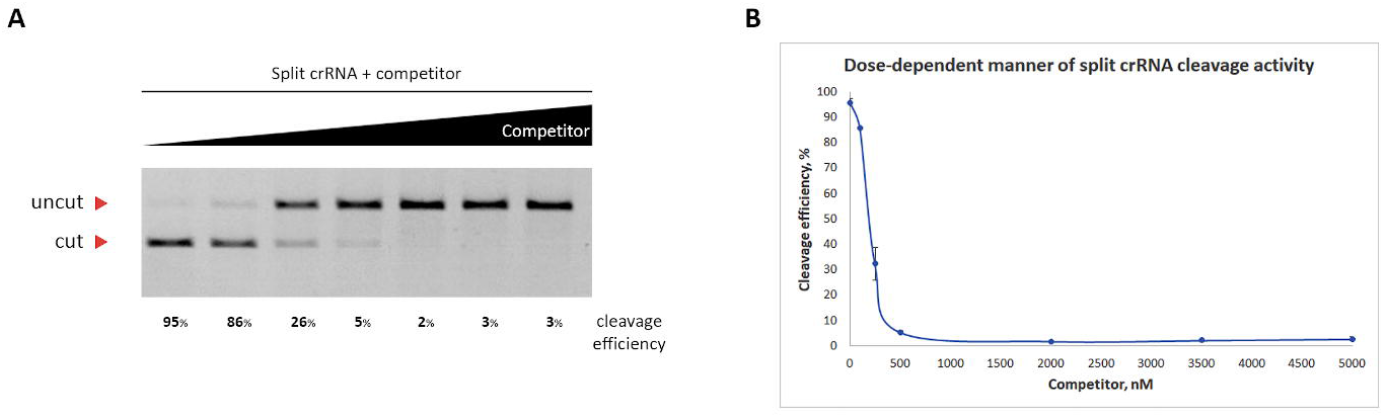
Effect of split crRNA on AsCas12a cleavage activity in presence of crRNA competitor. **(A)** The reactions contained 10 nM of PCR-amplified target DNA fragments, 500 nM of recombinant AsCas12a, and 5000 nM of split crRNAs, and increasing amounts of competitor crRNA (the ratios of split crRNA to crRNA competitor are 50:0, 50:1; 50:2,5; 50:5; 50:20; 50:35; 50:50), and were incubated 30 min at 37°C. **(B)** Quantification of the data presented on panel (A).

Given that the +2 extended scaffold version efficiently promoted target cleavage by spacer RNA (Figure 2C), we investigated the importance of the 5’- and 3’-extremities of the scaffold and spacer RNA domains for split crRNA function. Split versions bearing 5’- or 3’-terminal biotin (B) modifications on the spacer and scaffold RNAs were prepared and tested in the *in vitro* cleavage assay (Figure 5A). Among all the possible combinations of modified and unmodified RNAs, only the combination of 3’-modified scaffold RNA and 5’-modified spacer RNA significantly (∼50%) decreased the DNA substrate cleavage efficiency (Figure 5A), presumably because of steric interference at the scaffold and spacer RNA junctions caused by modifications. Indeed, when scaffold and spacer RNAs elongated by two non-complementary nucleotides towards the junction site were used to charge the effector, the cleavage activity was also decreased ∼50% at the conditions of the experiment (Figure 5B).

**Figure 5.**
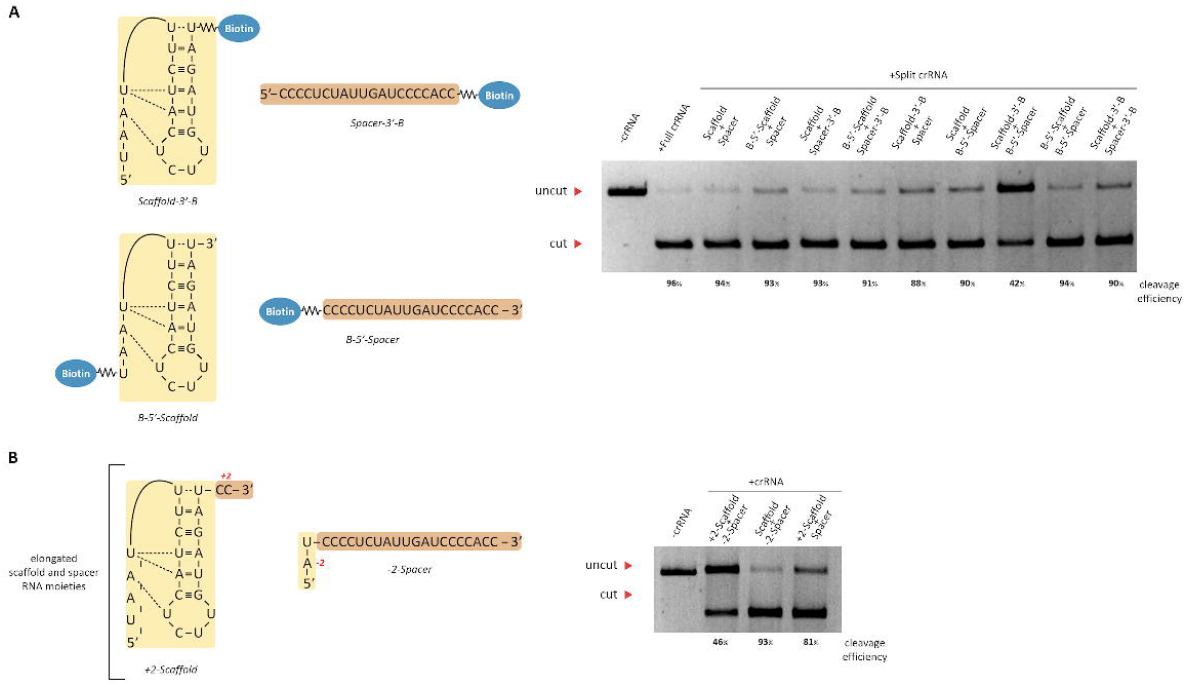
Effect of terminal modifications of spacer and scaffold RNAs on the activity of split crRNA. **(A)** Biotin modifications (shown as blue circles). **(B)** Two nucleotides elongated versions. The reactions contained 10 nM of PCR-amplified target DNA fragments, 500 nM of recombinant AsCas12a, and 5000 nM of crRNAs, and were incubated 30 min at 37°C.

### Target cleavage by AsCas12a programmed with spacer RNA only

Because split crRNA/AsCas12a cleaved the DNA template quite efficiently, we wondered whether the spacer domain alone could also induce cleavage. We surmised that though no cleavage was obtained in the presence of spacer RNA in the experiment shown in Figure 3, increasing the concentration of spacer RNA may allow the binding to the effector and target cleavage. An alternative scenario would be that the binding of the scaffold RNA is required for productive interaction with the spacer RNA by, for example, inducing a conformational change in the effector. Accordingly, various concentrations of spacer RNA as well as its 5’- and 3’-biotin modified versions were tested. Scaffold RNA was used as a control. As can be seen from Figure 6A, increasing the concentration of spacer RNA from 5000 to 15000 nM led to dose-dependent DNA substrate cleavage by AsCas12a. The reaction was strongly compromised by the presence of the 5’-end biotin modification. The 3’-end biotin modification did not affect the cleavage efficiency, as expected. Also as expected, no cleaved substrate was detected in presence of scaffold RNA alone, even at highest concentrations (Figure 6B).

**Figure 6.**
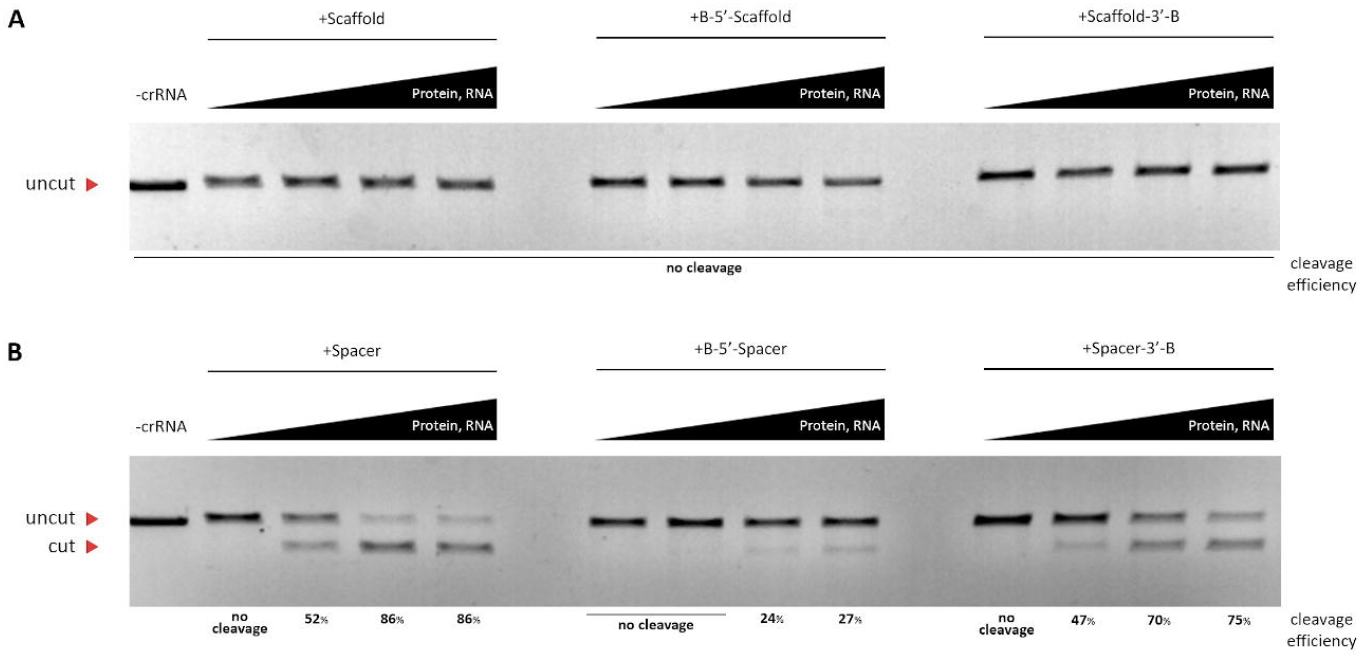
*In vitro* cleavage assay by purified AsCas12a in presence of increasing concentrations of scaffold **(A)** and spacer **(B)** parts of crRNA that were incubated 30 min. These concentrations correspond to 1:50:100, 1:50:500, 1:50:1000, 1:50:1500 DNA-AsCas12a-crRNA ratios, respectively.

### Analysis of target binding by the AsCas12a complex with split crRNA

To obtain further information on the split crRNA-AsCas12a complex and its interaction with target DNA, a fluorescence Cas beacon method was implemented, following a strategy that was earlier used to monitor the interaction of various CRISPR-Cas effector complexes with guide RNAs and target DNAs (12, 14, 15). Schematic representations of the beacon assay and structure of beacon used in this work are shown in Figures 7A and 7B. The PAM-distal ends of the beacon target and non-target strands are labeled with a fluorophore and a quencher, respectively. The baseline fluorescence intensity of a beacon is low due to quenching of the fluorescence label by the nearby quencher via FRET mechanism. Once incubated with a CRISPR-Cas effector complex, the beacon is recognized by the complex as a target DNA (Figure 7A). Binding of a CRISPR-Cas effector complex to beacon leads to separation of the fluorophore and the quencher and a readily detectable increase in fluorescence intensity (12, 14, 15).

**Figure 7.**
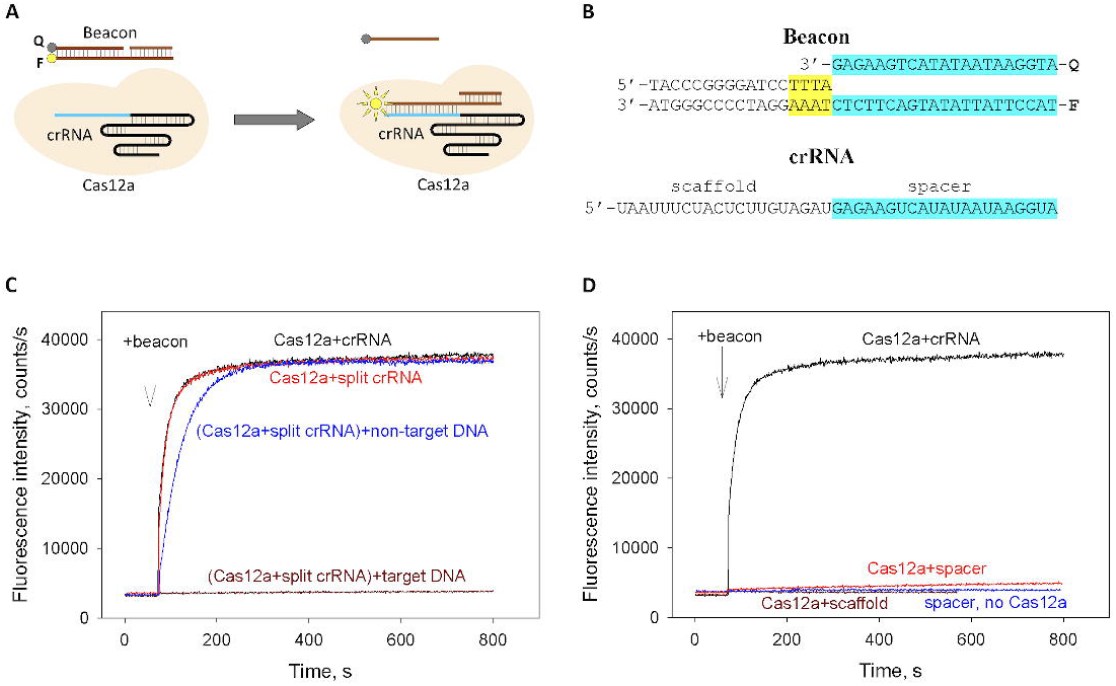
Cas beacon assay for AsCas12a interactions with intact crRNA, split crRNA and its scaffold and spacer fragments. **(A)** Schematic representation of beacon binding to crRNA-Cas12a. The circles labeled F and Q indicate the fluorophore and quencher. **(B)** Structures of crRNA and beacon. The crRNA spacer and beacon protospacer sequences are highlighted in blue, the PAM sequence is highlighted in yellow. **(C)** Time dependence of the increase in fluorescence upon the addition of 1 nM beacon to 40 nM AsCas12a combined with 60 nM of intact crRNA or split crRNA and inhibitory effects of target and non-target DNA competitors on beacon binding to split crRNA-AsCas12a. **(D)** Time dependence of the increase in fluorescence upon the addition of 1 nM beacon to samples containing 40 nM AsCas12a preincubated with 60 nM of crRNA, scaffold or spacer crRNA fragments and to a sample with 100 nM spacer in the absence of AsCas12a.

Upon the addition of beacon to 40 nM AsCas12a pre-incubated with 60 nM crRNA, fluorescence intensity rapidly increased, reaching peak intensity in about a minute (Figure 7C). Strikingly, the kinetics of beacon binding to AsCas12a preincubated with split crRNA was similar to that observed with intact crRNA (Figure 7C), thus implying that the AsCas12a complexes with intact and split crRNAs have comparable DNA binding activities. Next, we performed competition assays in which beacon binding to split crRNA-AsCas12a was monitored in the presence of either target DNA or non-target DNA that that bore no crRNA guide sequence complementarity. Preincubation of split crRNA-Cas12a with target DNA for 15 min led to severe inhibition of beacon binding, while non-target DNA exerted only a modest effect (Figure 7C). This finding indicated that split crRNA-AsCas12a formed a stable and specific complex with target DNA.

The fluorescence signal increased very slowly upon the beacon addition to sample containing 40 nM AsCas12a and 60 nM spacer RNA (Figure 7D). The rate of signal increase in this experiment was about 500-fold lower than that observed when AsCas12a was complexed with intact or split crRNAs. This observation suggests that spacer RNA alone confers AsCas12a only a low specific DNA binding activity in the absence of the scaffold segment. No increase in beacon fluorescence intensity was observed in the presence of spacer RNA without AsCas12a or in the presence of Cas12a preincubated with the scaffold RNA (Figure 7D).

### Split sgRNA/SpCas9 is not active *in vitro*

To determine whether the ability to use split RNAs is a common property of Class 2 effectors other than Type V AsCas12a, we performed *in vitro* target DNA cleavage assay using type II-A SpCas9 nuclease. The effector was charged with a single-guide RNA or its split version consisting of a 20-nt long spacer RNA complementary to the target and longer RNA corresponding to the rest of sgRNA (referred to as “Scaffold” in Figure 8). Only very low level of DNA cleavage by SpCas9 was detected even at the highest concentrations of split RNA components (Figure 8). It thus appears that unlike AsCas12a, SpCas9 is unable to efficiently use split crRNA for target recognition and cleavage.

**Figure 8.**
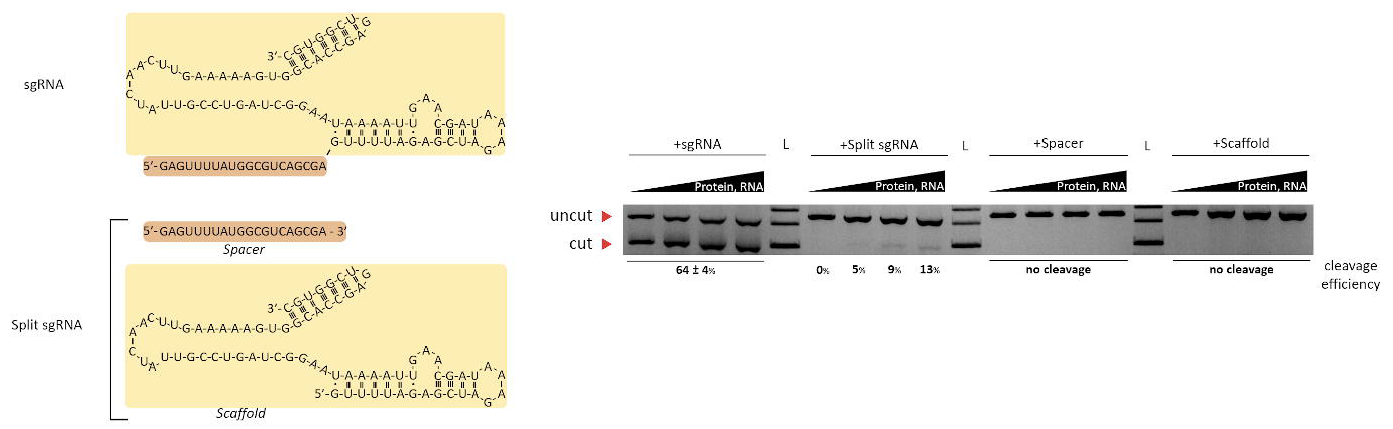
Effect of full-length and split sgRNA forms on SpCas9 cleavage activity. On the left, sequence of the sgRNA used in the study, its full-length and split forms. On the right, the products of *in vitro* cleavage of DNA template by purified SpCas9 in presence of increasing amounts (1000, 3000, 6000, and 8000 nM) of various sgRNA forms and SpCas9 (as indicated above panels).

### Split crRNA/AsCas12a is active in lysates of human cells

We next tested whether split crRNA can promote specific target cleavage in eukaryotic cell lysates. A human cell line expressing hAsCas12a in a Tet-inducible manner was constructed, and cleavage of target DNA in lysates of induced cells supplemented with various crRNAs -- intact, split, or individual spacer and scaffold -- was performed. The results, shown in (Figure 9) revealed hAsCas12a was indeed able to cleave the target as efficiently as intact crRNA. While spacer RNA alone was not effective, high (above 10000 nM) concentrations of this RNA also caused appreciable target cleavage in the lysate (data not shown).

**Figure 9.**
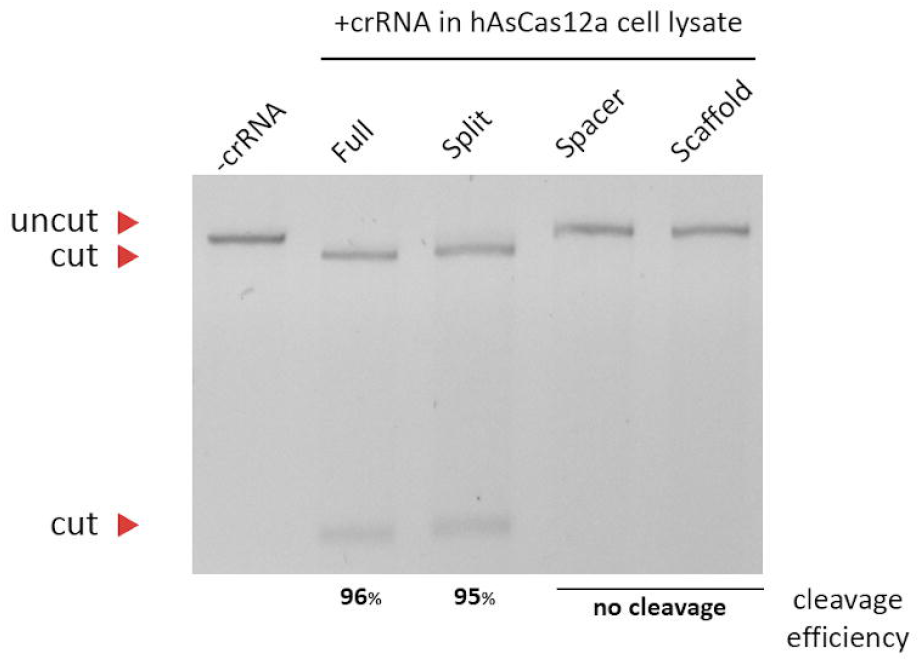
Cleavage of DNA template in lysates of human cells expressing hAsCas12a. The products of DNA cleavage in presence of various crRNA forms are shown. The cleavage efficiency is indicated below the panel as the ratio between cut and uncut DNA bands intensities. Cleavage reactions contained 4 nM of DNA template and 120 nM of crRNA. For the split crRNA variant, each component was added at the same concentration as the intact variant.

Taken together, our results demonstrate that class 2 Type V effector protein AsCas12a can use split crRNAs composed of two RNA molecules corresponding to the scaffold and spacer domains for specific cleavage of DNA template.

## DISCUSSION

Intact type V-A crRNA consists of two equally sized moieties, a scaffold (also referred to as direct repeat or 5’-handle) and a spacer (or guide) segment. Crystal structures of AsCas12a/crRNA complexes revealed that the crRNA scaffold adopts a pseudoknot structure and is tightly bound in a groove between two domains of AsCas12a in a sequence-specific manner (5). Comparison of crRNA structure in a binary Cas12a/crRNA and ternary Cas12a/crRNA/dsDNA complexes revealed that the scaffold pseudoknot structure is ordered in both complexes, while the entire spacer part is disordered in the binary complex (6). It has been suggested that the binding of Cas12a with PAM sequence in template DNA can induce conformational changes resulting in the pre-ordering of the crRNA seed sequence (nucleotides 1–8 of crRNA spacer) (4, 7). Next, the heteroduplex between the target DNA strand and the spacer part of crRNA is accommodated in the positively charged central channel between two lobes of the protein, in a sequence-independent manner.

The AsCas12a crRNA nucleotides U-10 and A-18 form a reverse Hoogsteen A:U base pair and participate in pseudoknot formation, which is additionally stabilized by non-canonical hydrogen bonds between U-10 and A-19 and between A-12, U-13, and U-17. The scaffold part of crRNA is tightly bound in the groove between two domains of Cas12a in a sequence-specific manner (5). Previous studies analysed the crRNA structure requirements for DNA cleavage by LbCas12a and FnCas12a nucleases (6, 7). In the case of FnCas12a, deletion of 5’-terminal scaffold nucleotides A-19, A-18 and U-17 had little effect on DNA cleavage efficiency (8). When 5’-terminal deletion also removed U-16, the DNA cleavage was seriously affected, and further shortening of the scaffold completely abolished cleavage. Since A-19, A-18 and U-17 are implicated in the pseudoknot formation, deletion of these three bases should turn the pseudoknot structure into a hairpin. U-16 forms a non-canonical pair with U-1, suggesting that the hairpin structure rather than the pseudoknot is required for efficient target recognition and cleavage. Indeed, mutations that disrupt the helical part of the hairpin abolish target cleavage by FnCas12a. In contrast to these findings, our analysis of the AsCas12a system shows, surprisingly, that deletions of large parts of the scaffold including residues participating in secondary structure formation have relatively minor effects on target DNA cleavage by AsCas12a. For example, we detected efficient target DNA cleavage when the crRNA scaffold part was shortened to just 7 bases, thus completely eliminating the 5’-helix structure. Detailed comparison of the intermolecular contacts in AsCas12a and FnCas12a-RNA-DNA ternary complexes (5, 7) reveals that in the AsCas12a complex, only the phosphodiester backbone of the crRNA scaffold helical part (except U-1 and U-16) provides the interactions with the protein, whereas, in the FnCas12a complex there are additional interactions of the G-3 and U-5 bases with the protein. This difference may explain the less stringent requirements of crRNA binding to AsCas12a compared to FnCas12a.

Given the results of structural analysis and our functional data, we considered whether the scaffold and spacer parts of crRNA can function separately. The 20-nt spacer RNA, when present in sufficiently high concentrations, induced target cleavage by AsCas12a. Thus, the spacer sequence alone can induce specific DNA cleavage by the nuclease. For the LbCas12a system, it was demonstrated that removal of the spacer sequence abolished DNA cleavage but only partially compromised the affinity between the crRNA and the effector. Furthermore, an excess of crRNA scaffold inhibited DNA cleavage by LbCas12a charged with intact crRNA (6), demonstrating that the scaffold can compete with intact crRNA for the binding to LbCas12a. Our data also show that crRNA scaffold alone binds to AsCas12a. Most significantly, the scaffold, when present separately, strongly increased the ability of separated spacer RNA to induce target cleavage by AsCas12a. Furthermore, our data show that AsCas12a combined with separated spacer and scaffold forms a stable complex with target DNA. The functionality of such split crRNA is the principal finding of our work. The result indicates that the scaffold not only interacts with AsCas12a but also increases the efficiency of proper binding of the spacer in either the binary, ternary (or both) complexes, presumably by inducing a conformational change in the effector.

The functionality of split crRNA system may help further development of genome editing applications using AsCas12a. For example, co-expression of the scaffold RNA together with several spacer sequences might be useful for facile multiplexing of editing protocols. Further, since both split crRNA components can withstand additional modifications without the loss of function, targeting specific sites, including organelles surrounded by the double membrane (16) may become possible. Studies from several laboratories demonstrated that while long non-coding RNAs (75-300 nucleotides) can be imported into mitochondria with the help of specific protein factors (17), shorter RNA molecules such as miRNA and siRNA are targeted in a much more efficient and non-specific way (18). By reducing the lengths of crRNA components, the split crRNA/Cas12a system may thus be targeted into human mitochondria and allow efficient mitochondrial genome editing.

